# Large language models overcome the challenges of unstructured text data in ecology

**DOI:** 10.1101/2024.01.23.576654

**Authors:** Andry Castro, João Pinto, Luís Reino, Pavel Pipek, César Capinha

**Affiliations:** Centro de Estudos Geográficos, Instituto de Geografia e Ordenamento do Território, Universidade de Lisboa, Lisboa, Portugal; Global Health and Tropical Medicine, Instituto de Higiene e Medicina Tropical, Universidade Nova de Lisboa, Lisboa, Portugal; CIBIO, Centro de Investigação em Biodiversidade e Recursos Genéticos, InBIO Laboratório Associado, Campus de Vairão, Universidade do Porto, Vairão, Portugal; BIOPOLIS Program in Genomics, Biodiversity and Land Planning, CIBIO, Vairão, Portugal; CIBIO, Centro de Investigação em Biodiversidade e Recursos Genéticos, InBIO Laboratório Associado, Instituto Superior de Agronomia, Universidade de Lisboa, Lisboa, Portugal; Department of Invasion Ecology, Institute of Botany, Czech Academy of Sciences, Průhonice, Czech Republic; Department of Ecology, Faculty of Science, Charles University, Prague, Czech Republic; Associate Laboratory Terra, Portugal

**Keywords:** AI, Automation, Data integration, GPT, LLaMA, Unstructured data

## Abstract

The vast volume of currently available unstructured text data, such as research papers, news, and technical report data, shows great potential for ecological research. However, manual processing of such data is labour-intensive, posing a significant challenge. In this study, we aimed to assess the application of three state-of-the-art prompt-based large language models (LLMs), GPT 3.5, GPT 4, and LLaMA-2-70B, to automate the identification, interpretation, extraction, and structuring of relevant ecological information from unstructured textual sources. We focused on species distribution data from two sources: news outlets and research papers. We assessed the LLMs for four key tasks: classification of documents with species distribution data, identification of regions where species are recorded, generation of geographical coordinates for these regions, and supply of results in a structured format. GPT 4 consistently outperformed the other models, demonstrating a high capacity to interpret textual data and extract relevant information, with the percentage of correct outputs often exceeding 90% (average accuracy across tasks: 87–100%). Its performance also depended on the data source type and task, with better results achieved with news reports, in the identification of regions with species reports and presentation of structured output. Its predecessor, GPT 3.5, exhibited reasonably low accuracy across all tasks and data sources (average accuracy across tasks: 81–97%), whereas LLaMA-2-70B showed the worst performance (37– 73%). These results demonstrate the potential benefit of integrating prompt-based LLMs into ecological data assimilation workflows as essential tools to efficiently process large volumes of textual data.

## 1. Introduction

Recently developed conversational artificial intelligence (AI)-based large language models (LLMs), such as GPT-4 by OpenAI (Brown et al., 2020), LLaMA series by Meta (Touvron et al., 2023ab), have attracted significant attention. The transformer architecture underlying these models, particularly their self-attention mechanism, enables them to handle complex large-scale data efficiently and capture intricate patterns within it (Devlin et al., 2018). These models show exceptional ability to interpret user requests, provide sensible responses, and demonstrate proficiency in various skilled tasks involving the analysis and processing of digital data (Bommasani et al., 2021). Their applications are vast, ranging from natural language understanding to more specialised tasks, such as code generation and multimodal interactions, where models process and generate text based on image inputs. These models can aid in the advancement of various scientific fields, including ecology, by enabling automated data analysis, improving the accuracy of ecological modelling, and facilitating the interpretation of complex datasets (Morera, 2024).

One area that can greatly benefit from the advancements facilitated by LLMs is information extraction and structuring, particularly from textual data. The large amount of unstructured textual data from various sources, including research papers, online information on social media, and news articles, poses a significant challenge for ecologists. Such information is invaluable for both basic and applied ecological research, supporting many applications, including sentiment analysis of exotic pets (Moloney et al., 2021), quantification of temporal ecological patterns (Hart et al., 2018), and comprehensive mapping of species distribution (Monteiro et al., 2020). However, the unstructured nature of textual data necessitates several challenging information search and processing steps, including distinguishing between the relevant and irrelevant data sources, interpreting the textual information, and converting it into structured formats suitable for analysis. Although various tools have been proposed to facilitate these tasks (Cornford et al., 2020; Le Guillarme and Thuiller, 2022), they mostly involve extremely labour-intensive and time-consuming manual procedures, which severely limit the continuous and large-scale analysis of textual data.

Considering the importance of these tasks, several studies have assessed the capacity of computational models to process textual ecological information. The models used range from classical machine learning algorithms, such as Naive Bayes (Stringham et al., 2021), to more recent transformer-based models trained on domain-specific datasets, which is a typical requirement to ensure high performance (Lee et al., 2020). However, only a few studies have examined the performance of new conversational or prompt-based models, such as GPT and LLaMA. These models, pretrained on extensive multidomain datasets, can perform tasks based on simple text-based instructions provided by users (prompts), thus eliminating the need for resource-intensive and complex domain-specific model training. Recent studies using prompt-based approaches include those by Gougherty and Clipp (2024), who tested the text-bison-001 generative text model of Google to extract information from 100 disease reports, and Scheepens et al. (2024), who evaluated the ability of GPT-4 to extract information on invertebrate pests and pest controllers from research paper abstracts. Both studies underscored the potential of prompt-based LLM approaches. However, to date, no studies have comprehensively assessed the performances of various prompt-based LLMs across the three main steps of information extraction and structuring from text data: document classification (i.e. determining if the source contains relevant information), information extraction, and data structuring as per the predefined data formats. Additionally, owing to their diversity, the performance of different LLMs may vary across tasks. Therefore, comparative evaluation of these models is necessary to determine their relative effectiveness in extracting and structuring ecological information.

In this study, we evaluated the abilities of multiple prompt-based LLMs to identify, extract, and structure relevant ecological information from textual data. We focused on species distribution data, which is crucial for assessing biodiversity change (Boonman et al., 2024), conservation needs (Guisan et al., 2013) and mapping the spread of invasive alien species (Latombe et al., 2017). Currently, these data are available in textual form in various sources, including scientific literature (Maquart et al., 2021), technical reports (Mota et al., 2006), and multiple online platforms, such as news websites, social media sites (Chowdhury et al., 2023), and institutional webpages. Owing to the inherent difficulty of manually handling these data, researchers often overlook these sources, leading to distributional information gaps that hinder research and decision-making in conservation and environmental management. The rapidly growing volume of scientific and non-scientific publications (Landhuis, 2016) and low adoption of structured data reporting standards (Castro et al., 2023; Poisot et al., 2019) further exacerbate the information gap.

Here, we measured and compared the performances of three popular state-of-the-art prompt-based LLMs, including two commercial models (GPT 3.5 and GPT 4; OpenAI, 2023) and one open-source free-to-use model (LLaMA-2-70B; Touvron et al., 2023ab). We evaluated these models across four key tasks in the automated extraction of ecological information from textual data: *i*) ability to distinguish relevant from irrelevant data sources, *ii*) accuracy of the information extracted, *iii*) capability to geocode spatial entities mentioned in the text, and *iv*) provision of the requested information in a structured format. The assessment was performed using two types of textual data sources: research papers and online news articles, on two globally important invasive species.

## 2. Material and Methods

Here, we simulated the applications of LLMs in real-world settings and used them to process the unstructured textual data from news outlets and research papers on two range-expanding problematic invasive species. We aimed to extract the distribution data from these sources to better track the ongoing range expansion of invasive species. We evaluated four key tasks ranging from information identification to integration, including the ability of the models to 1) identify research papers and news articles in different media outlets providing species observation data, 2) extract the names of regions with reported observations, 3) generate geographical coordinates for the identified regions, and 4) present the information in a consistent structured format.

### 2.1 Establishing a test dataset

We used the Asian hornet (*Vespa velutina*) and Asian tiger mosquito (*Aedes albopictus*) as test cases in our study. These species are highly problematic in their alien range, causing economic damage and threatening human health (Barbet-Massin et al., 2020; Schaffner et al., 2013), and spreading at a fast rate to new regions, particularly in Europe (Monceau and Thiery, 2016; Schaffner and Mathis, 2014). Therefore, there is a significant interest in researchers and entities tasked with invasion prevention strategies to track the geographical spread of these species. However, a relevant volume of distributional information is provided in the form of textual narratives delivered in news (Bullens, 2023; Carballo, 2023) or research papers (Bakran-Lebl et al., 2021; Dillane et al., 2022).

As a representative of the current unstructured textual data on the distribution of these species, we considered two sources: research papers and online newspapers. Researchers have published a relevant number of works on these species. However, only a limited number of them tend to provide distributional data, with another common focus being health or economic impact-related information and genetic studies. Similarly, online newspaper news provides key and often timely information on new species observations (Vuorisalo et al., 2001). However, this is often mixed with news on several other subjects, even if related to the same species, such as precautions to be taken and the potential economic or health impacts. Manual sorting through this vast and diverse content to identify reports of new species observations is not only challenging, but also often unsustainable because of the sheer volume and complexity of the data.

To incorporate these two types of data source, we used two programmatic interfaces for the R Language (R Core Team, 2022): the ‘openalexR’ (Aria et al., 2023) and the ‘newsanchor’ (Frie et al., 2019) packages. The first is an R wrapper for the Open Alex API (https://openalex.org) which enables users to query and extract data from the Open Alex database which includes over 250M academic works. It provides access to multiple fields of the work including titles, abstracts, and publication dates. The ‘newsanchor’ package connects to the News API (https://newsapi.org), allowing it to search and retrieve live news headlines from over 30,000 news sources and blogs. Here, we used the developer licence of the News API, which has some limitations, including searching only for news up to one month old. Crucially, both packages allow near-real-time access to published information, making them suitable for the continuous and timely monitoring of new species distribution data.

We searched for research papers and news articles using scientific names and common names of species in English in the case of research papers and in ten different languages for the news (Dutch, English, French, Italian, German, Norwegian, Portuguese, Russian, Spanish, and Swedish; e.g. *A. albopictus*, tiger mosquito, zanzara tigre, and asiatisk tigermygg). For research papers, we saved titles, abstracts, and DOIs. In several cases, the abstracts were not available from ‘openalexR’, so we obtained these using the ‘rcrossref’ package (Chamberlain et al., 2022). For news articles, we saved the title and description (highlight). Owing to limitations in API access and unequal volumes of results, the period used in the searches differed between species and source types. In each case, the temporal extent preceding the date of search retrieval was extended until 150 results were obtained. This resulted in a dataset of 600 records for testing (2 × 150 records for each species; **Table 1**). For each of these records, we manually identified whether it reported the species’ distributional information, and if so, for which region or regions (**Table 1** and see the full database of news and papers in **Appendix A** and the R code for news and paper retrieving in **Appendix B**).

**Table 1.**
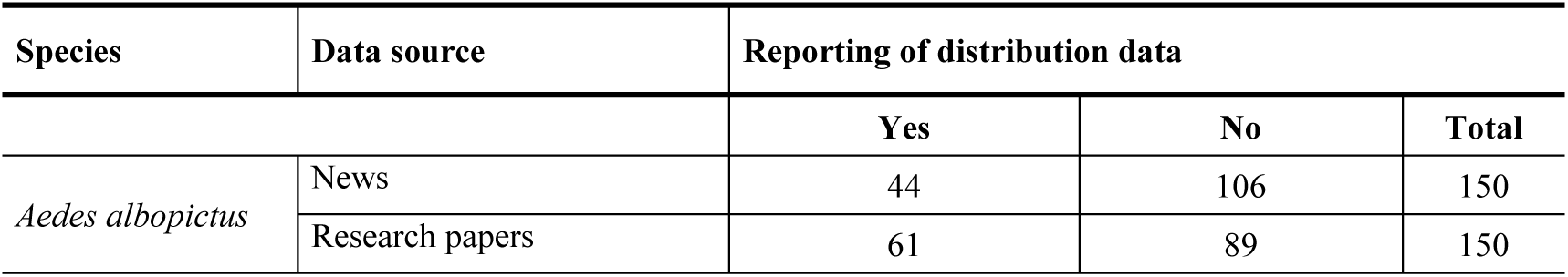
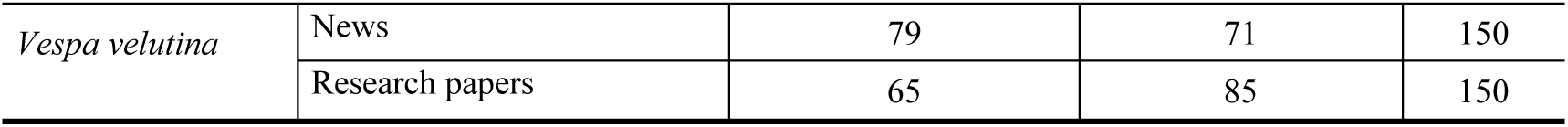
Composition of the dataset assembled to test the capacities of large language models to identify data distribution and extract information for two invasive species: *Aedes albopictus* mosquito and *Vespa velutina* hornet.

| Species | Data source | Reporting of distribution data |  |  |
| --- | --- | --- | --- | --- |
|  |  | Yes | No | Total |
| *Aedes albopictus* | News | 44 | 106 | 150 |
|  | Research papers | 61 | 89 | 150 |
| *Vespa velutina* | News | 79 | 71 | 150 |
|  | Research papers | 65 | 85 | 150 |

### 2.2 Prompt-based testing approach

We implemented a few-shot learning approach (Brown et al., 2020; Chiu et al., 2021), providing LLMs examples of data sources with or without reported species distributions. We performed this separately for the research papers and news articles. For the former, we provided a set of eight positive and eight negative cases, each characterised by a title and abstract. For news, we used the same procedure, giving the news title alongside the description (highlight) of the news. In some cases, however, we provided only titles because news descriptions or paper abstracts were not always provided by the search results.

Then, we instructed the models to generate a simple answer of “YES” or “NO” to identify if papers or news to be examined include or not (respectively) distribution data of the species. For positive classifications, we also instructed the model to provide the name or names of the regions where the species was mentioned and, in the following line, to provide its geographical coordinates in decimal degrees. We also provided models with examples of the expected results in both cases. Additional species-relevant instructions were also provided. Specifically, we informed LLMs about possible confusions between species common names (e.g., Asian hornet vs Asian giant hornet, the latter being a distinct species) and to exclude research papers related to potential (i.e., not actually observed) distributions and papers or news referring to geographical resolutions coarser than that of a single country (e.g., a set of two or more countries, or a continent). Finally, we also asked the model to provide the results in a consistent, standardised, format, having the classification result (i.e., ‘YES’ or ‘NO’) in the first line, and for positive cases the region name(s) in the second line and the coordinates in the third (last) line (See the full text prompt in the **Appendix C**).

### 2.3 Data testing

We performed our tests using three recent large-language models: GPT-3.5, GPT-4 from OpenAI (OpenAI, 2023) and LLaMA-2-70B (Touvron et al., 2023ab) from Meta-AI. The Generative Pretrained Transformer (GPT) series, which is based on the transformer architecture introduced by Vaswani et al. (2017), represents a major advancement in the field of LLMs. The GPT-3 model presented by Brown et al. (2020) features an autoregressive LM with 175 billion parameters, trained on a vast text corpus. Its enhanced version, GPT-3.5, incorporates reinforcement learning from human feedback (Ouyang et al., 2022) to improve performance, and is currently available in a limited chat mode from OpenAI. The subsequent model, GPT-4, is larger and more advanced, although its exact parameter count remains unknown. The use of GPT-4 is restricted to commercial licences and is considered the current state-of-the-art method for large LLMs. LLaMA-2-70B, an open-source LLM, was pretrained on two trillion tokens of data, including instruction datasets and human-annotated examples. This model outperforms other open-source LLM models on most benchmarks (Touvron et al., 2023ab) and is a suitable substitute for closed-source models such as those from the GPT family. The models were tested using the web interface, i.e., chat.openai.com for the first two (GPT) models and llama2.ai for the LLaMA-2-70B model.

For each model, we performed 600 individual tests in a new dialog window by manually copying and pasting each prompt. Although the model did not receive any feedback on the correctness of the previous answers, we used this procedure to ensure that no other information from the previous requests influenced the response of the model to subsequent prompts. For each test, we evaluated 1) the correct classification of the news or paper in terms of the provision of distribution data, 2) the correct naming of regions for which distribution records were reported, 3) the provision of geographical coordinates falling within the region for which the distribution was reported, and 4) the provision of results in a specified structured format (the expected format is shown in Table 2 for sources reporting and not reporting a species). All tests were made from 11th of December 2023 to the 3rd of January 2024.

**Table 2.** Examples of the results obtained using the models in terms of classification of documents, correct identification of regions for which the species distribution is mentioned, supply of coordinates within the mentioned region, and provision of all results in the requested format. Two news articles reporting the presence of *V. velutina* in Jersey Island and a single news article on *A. albopictus* are provided. The first two correspond to positive cases (species distribution is reported), whereas the latter corresponds to a negative case, for which it is not possible to establish a relationship between the species and specific geographical area. Examples of incorrect and correct classifications and output generated by the models are provided in Appendices D and Appendix E, respectively.

| Examples<br>to classify | Expected<br>result | Parameters classified |  |  |  |
| --- | --- | --- | --- | --- | --- |
|  |  | Document<br>classification | Region<br>name | Coordinates | Format |
| Example of an expected "YES", directly reported |  |  |  |  |  |
| *"Invasive hornet species found in the Jersey Island for the first time - This marks the first time the yellow-legged hornets have been reported on the Island."* | "YES<br>Jersey Island<br>49.214439°, -2.131250°" | Correct | Correct | Correct | Correct |
| Example of an expected "YES", indirectly reported |  |  |  |  |  |
| *"Horriifying moment Asian hornet devours wasp in a Jersey Island school in just seconds -The video shows the moment that an Asian hornet devoured a wasp in seconds"* | "YES<br>Jersey Island<br>49.214439°, -2.131250°" | Correct | Correct | Correct | Correct |
| **Example of an expected "NO"** |  |  |  |  |  |
| *"Aedes albopictus is growing in urban areas of developed countries - Cities are facing a threat under new climatic conditions"* | "NO" | Correct | NA | NA | Correct |

We assessed model performance by measuring the accuracy (percentage of correct results) for each evaluated task and complementary true positive rate (TPR, also known as recall or sensitivity) and true negative rate (TNR, also known as specificity). For geographical coordinates, we compared the performances of our models with that of the current state-of-the-art approach, Nominatim 4.3.2 API (https://nominatim.org/release-docs/latest/api/), an open-source geocoding service provided by the OpenStreetMap (OSM) project, via R.

Examples of these model results are presented in **Table 2**. To facilitate the interpretation of model performance, accuracy levels are qualitatively classified as perfect (100%), very good (<100% and ≥ 90%), good (<90% and ≥ 70%), moderate (<70% and ≥ 50%) and poor (<50%).

## 3. Results

Overall, GPT-4 generally outperformed GPT-3.5, and both consistently outperformed LLaMA-2-70B across the four tasks assessed in this study **(Fig. 1)**.

**Fig. 1.**
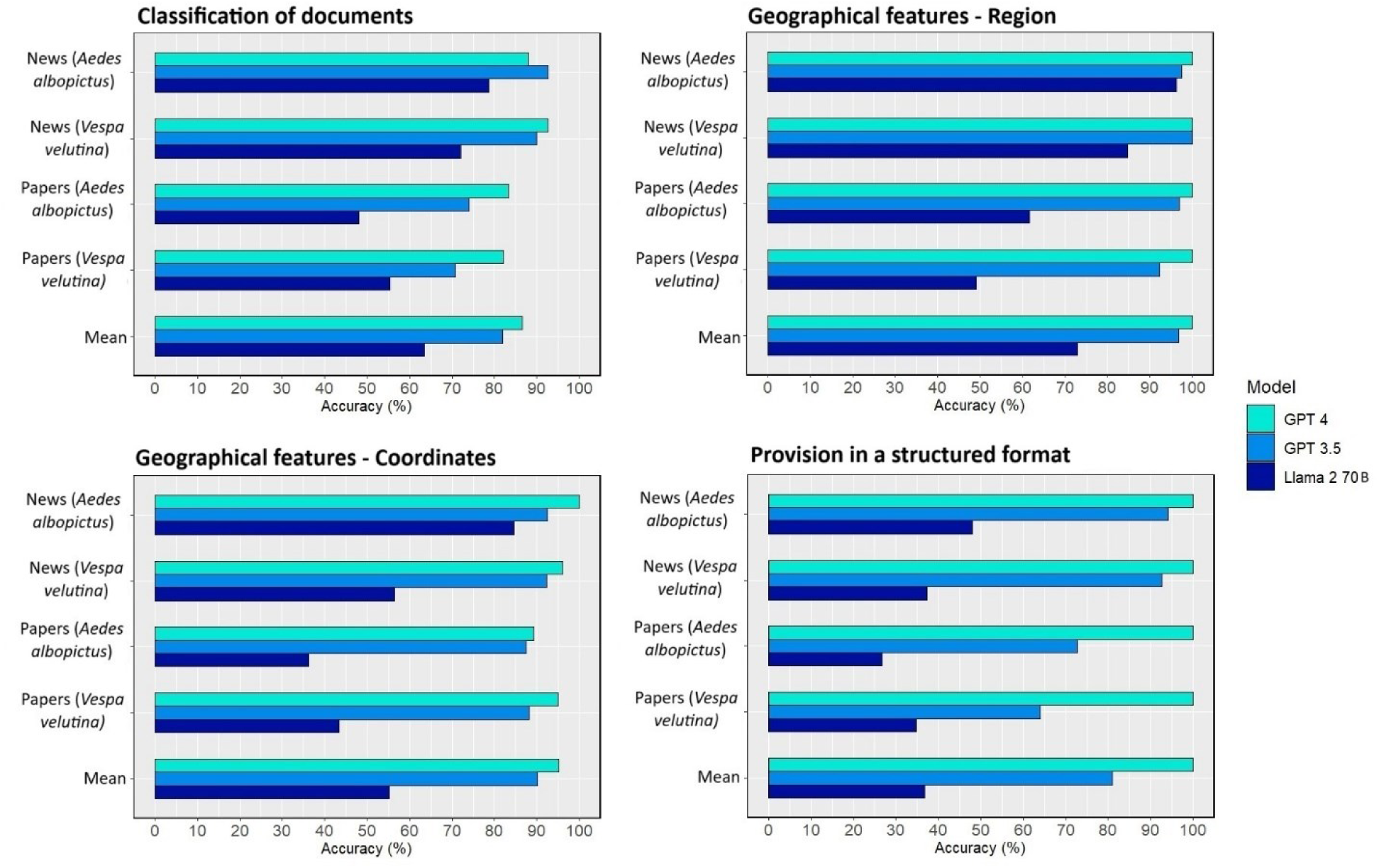
Accuracies of the tested models in (a) distinguishing between documents reporting the presence or absence of a given species, (b) providing the name(s) of region(s) with the specific species, (c) supplying coordinates falling in the region(s) identified, and (d) providing results in a structured format.

### 3.1 Classification of documents

The first task assessed the capacity of the models to identify unstructured data sources reporting the presence of a species **(Fig. 1a)**. GPT-4 exhibited good and very good levels of accuracy for news (*A. albopictus* = 88.0%; *V. velutina* = 92.7%) and good level of accuracy for papers (*A. albopictus* = 83.3%; *V. velutina* = 82.0%). GPT-4 is followed by GPT-3.5 with very good levels of accuracy for news (*A. albopictus* = 92.7%; *V. velutina* = 90.0%) and good levels of accuracy for papers (*A. albopictus* = 74.0%; *V. velutina* = 70.7%). LLaMA-2-70B had the least accurate results for this task, showing good levels of accuracy for news (*A. albopictus* = 78.7%; *V. velutina* = 72.0%) and poor-to-moderate levels of accuracy for papers (*A. albopictus* = 48.0%; *V. velutina* = 55.3%). Notably, GPT-4 was more accurate in identifying documents not reporting any species (TNR = 94.7% ± 4.6) than in identifying those reporting a specific species (TPR = 73.4% ± 16.6). In contrast, GPT-3.5 showed a smaller average difference, performing worse than GPT-4 in identifying negatives but better than GPT-4 in identifying positives (TNR = 81.9% ± 12.4 vs. TPR = 79.9% ± 19.9). The LLaMA-2-70B model exhibited the opposite trend, showing more accuracy in identifying positives (TPR = 69.9% ± 12.0) than in identifying negatives (TNR = 54.4% ± 32.1), but had markedly lower overall performance than the other models **(Table 3)**.

**Table 3.** True positive (TPR) and negative (TNR) rates of models classifying documents reporting the presence or absence of a species.

|  |  | Large language models classification |  |  |  |  |  |
| --- | --- | --- | --- | --- | --- | --- | --- |
|  |  | GPT-4 |  | GPT-3.5 |  | LLaMA-2-70B |  |
| Specie name | Type of document | TPR (%) | TNR (%) | TPR (%) | TNR (%) | TPR (%) | TNR (%) |
| *Aedes albopictus* | News | 75.0 | 93.4 | 90.9 | 93.4 | 59.1 | 86.8 |
|  | Papers | 60.7 | 98.9 | 52.5 | 87.6 | 77.0 | 28.1 |
| *Vespa velutina* | News | 96.2 | 88.7 | 97.5 | 81.7 | 58.2 | 87.3 |
|  | Papers | 61.5 | 97.6 | 78.5 | 64.7 | 81.5 | 35.3 |
| Mean | | 73.35 ± 16.6 | 94.7 ± 4.6 | 79.9 ± 19.9 | 81.9 ± 12.4 | 69.0 ± 12.0 | 54.4 ± 32.1 |

### 3.2 Geographical features

The second and third tasks only considered news and papers reporting a specific species. Considering the naming of the region **(Fig. 1b)**, GPT-4 exhibited perfect levels of accuracy for news (*A. albopictus* = 100%; *V. velutina* = 100%) and papers (*A. albopictus* = 100%; *V. velutina* = 100%). GPT-3.5 achieved very good-to-perfect levels of accuracy for news (*A. albopictus* = 97.5%; *V. velutina* = 100%) and very good levels of accuracy for papers (*A. albopictus* = 96.9%; *V. velutina* = 92.2%). LLaMA-2-70B was the weakest model, exhibiting very good and good levels of accuracy for news (*A. albopictus* = 95.2%; *V. velutina* = 84.8%) and moderate and poor levels of accuracy for papers (*A. albopictus*= 61.7%; *V. velutina* = 49.1%).

Considering its capacity to provide coordinates within the identified regions **(Fig. 1c)**, GPT-4 exhibited perfect and very good levels of accuracy for news (*A. albopictus* = 100%; *V. velutina* = 96.1%) and good and very good levels of accuracy for papers (*A. albopictus* = 89.2%; *V. velutina* = 95%). GPT-3.5 exhibited very good levels of accuracy for news (*A. albopictus* = 92.5%; *V. velutina* = 92.2%) and good levels of accuracy for papers (*A. albopictus* = 87.5%; *V. velutina* = 88.2%). LLaMA-2-70B exhibited the lowest accuracy levels for this task, with good and moderate levels of accuracy for news (*A. albopictus* = 84.6%; *V. velutina* = 56.5%) and poor levels of accuracy for papers (*A. albopictus* = 36.2%; *V. velutina* = 43.4%).

Notably, GPT-4 results were generally comparable to those of the current state-of-the-art geocoding method, Nominatim. For the positive cases where all other tasks were correctly assessed, GPT-4 generated coordinates of the identified regions with an accuracy of 95.2% (177 correct coordinates out of 186 regions), which was comparable to that of Nominatim (96.2%; 179 correct coordinates out of 186 regions).

### 3.3 Provision of results in a structured format

Concerning the supply of results in the requested structure **(Fig. 1d)**, GPT-4 exhibited 100% accuracy for both data source types. GPT-3.5 achieved very good and very good levels of accuracy for news (*A. albopictus* = 94%; *V. velutina* = 92.7%) and good-to-moderate levels of accuracy for papers (*A. albopictus* = 72.7%; *V. velutina* = 64%). LLaMA-2-70B model showed the worst performance, with poor levels of accuracy for both news (*A. albopictus* = 48%; *V. velutina* = 37.3%) and papers (*A. albopictus* = 26.7%; *V. velutina* = 34.7%).

## 4. Discussion

In this study, we measured and compared the abilities of three new LLM models, GPT 3.5, GPT 4, and LLaMA-2-70B, to identify, interpret, and structure relevant ecological information from unstructured textual data sources. Our results revealed the potential of these models to accurately perform these tasks. However, their performance varied substantially based on the type of data source and specific LLM used.

Among the three compared LLMs, GPT-4 generally achieved the highest performance across the four evaluated tasks: classification of documents reporting distribution data, identification of regions with species reports, generation of geographical coordinates for these regions, and delivery of results in a structured format. The superiority of GPT-4 over the other models was expected as it is considered the state-of-the-art LMM and an upgrade over GPT-3.5, which showed improvements in several domains, including natural language interpretation (Espejel et al., 2023). It is important to note, however, that for certain types of input and tasks, particularly in the provision structured output, GPT-3.5 performed only slightly less effectively. Additionally, GPT models consistently outperformed LLaMA-2-70B in accuracy in all four tasks, often by over 40%. This large improvement margin suggests that LLaMA-2-70B is not an out-of-the-box solution for processing unstructured textual ecological data.

We also noted substantial differences in the models’ capacity to handle text from news articles versus research articles. Across the models, text classification, region extraction (the two tasks are directly dependent on the provided text), and the supply of results in a structured format (except GPT-4, where no difference was found) consistently showed better results for news-derived text than for papers. This is likely because of the more complex and convoluted narratives often found in research papers, which make model interpretation more challenging. In contrast, news typically aims to provide easily understandable text using simpler language. This difference in comprehension capacity between research papers and news is less pronounced in the GPT-4, especially in geographic region identification, further underscoring the need for large and complex models to understand scientific languages. Additional factors may also have played a role. Parameters such as the informational content of the examples in the prompt, the language of the text, or even the order in which the examples are provided could have an impact (Zhao et al., 2021). Although exploring the precise factors that drive the observed differences is outside the scope of this study, it represents a relevant avenue for future research.

Due to the use of distinct test sets, drawing direct comparisons between our results and those of previous studies is challenging. However, the models tested show comparable performance with the current state-of-the-art approaches. For instance, recent studies using custom-trained BERT models (a type of transformer architecture) to classify online texts of ecological relevance reported accuracies of 87–97% (Edwards et al., 2022; Hunter et al., 2023). Here, performance of the GPT models for news fell within this range, without the need for specific model training. Additionally, our findings substantiate those of Scheepens et al. (2024), who reported the good performance of GPT-4 in information extraction. Our results complement these findings by showing that GPT-3.5 also performs well in this task, with only a slight decrease in performance compared to GPT-4. Similarly, our results corroborate the findings of Gougherty and Clipp (2024), who documented the capacity of generative models to generate (i.e. not extract) accurate geospatial coordinates. We found this capacity of GPT-4 to be comparable to that of Nominatim, the current standard in automated geocoding (e.g., Otero et al., 2024). Finally, this study is among the first to assess the ability of generative AI models to provide responses in a structured format, which is a critical requirement for integration into automated workflows. This aspect, which has not been extensively addressed in previous studies, can serve as a benchmark for future research in this area.

Overall, capabilities of the tested GPT models, particularly GPT-4, were remarkable. Integrating GPT models into data integration workflows will significantly reduce the manual workload and offer a promising avenue to process the growing volume of unstructured ecological data. The programmatic integration of such models can be facilitated by specifically developed tools and libraries for programming languages commonly used by ecologists, such as R (Rodriguez, 2023) and Python (Arunachalam, 2023). We anticipate that these tools will become increasingly available over time. Currently, various tools are available for the automated parsing of unstructured text data that are essential for effectively feeding these models. Although we focused on processing short-text sequences from news articles and scientific papers, which are crucial for near-real-time monitoring of new species occurrences (e.g., Capinha et al., 2023), these models can also be used to handle large and complex documents, such as full research papers in PDF format. Fortunately, tools now exist that deliver high-quality deparsed text, ready for interpretation in LLMs (e.g., DocParser; Rausch et al., 2021).

Nonetheless, several challenges persist in the use of GPT models, especially in terms of the financial resources required for implementation. Although processing a typical research paper using GPT-4 is relatively affordable (< $1) (OpenAI, 2024), scaling up thousands of papers can impose significant financial constraints. Programmatic licences and usage caps also have limitations. Open-source LLMs, such as LLaMA-2-70B, which are relatively less accurate, can be used as alternatives after necessary fine-tuning (Wang et al., 2024). These open-source LLMs can overcome some limitations of commercial models, particularly regarding API access and usage limits. Technological advancements can further improve the open-source LLM performance. However, operating these models locally, particularly for larger variants, requires substantial computational resources, including high-end graphic processing units (Kodali et al., 2024), further increasing the acquisition, operational, and maintenance costs.

## Conclusion

In conclusion, this study highlights the remarkable ability of next-generation prompt-based LLMs to assimilate and process unstructured textual data and deliver them in a structured format suitable for ecological data processing. However, the models exhibited variable performance depending on the data source and specific task. Therefore, further comparative testing is necessary to identify the most effective settings. Despite the implementation challenges, integration of LLMs will create seamless workflows for the analyses and processing of unstructured textual information, particularly that on species distribution information, in ecological research.

## Author contributions

AC and CC conceived the study. AC and PP performed the analyses. AC led manuscript writing with CC. All authors contributed to writing and editing.

## Data availability statement

The data used in this study is provided in the appendices.

## Acknowledgements

The authors acknowledge the constructive suggestions from three reviewers that helped improved this work.

## Abbreviations

LLMs: large language models
TPR: true positive rate
TNR: true negative rate

**Appendix A.** Database of news and papers.

**Appendix B.** R code for news and papers scraping and R code used for the geocoding.

**Appendix C.** Full text prompt.

**Appendix D.** Examples of incorrect classifications by the LLM’s.

**Appendix E.** Correct classifications by the LLM’s.

